# Micrococcal nuclease regulates biofilm formation and dispersal in methicillin-resistant *Staphylococcus aureus* USA300

**DOI:** 10.1101/2023.11.05.565664

**Authors:** Jeffrey B. Kaplan, Aleksandr P. Florjanczyk, Maria Ochiai, Caleb D. Jones, Alexander R. Horswill

## Abstract

Biofilm formation is an important virulence factor for methicillin-resistant *Staphylococcus aureus* (MRSA). The extracellular matrix of MRSA biofilms contains significant amounts of double-stranded DNA. MRSA cells also secrete micrococcal nuclease (Nuc1) which degrades double-stranded DNA. In this study we used a *nuc1* mutant strain to investigate the role of Nuc1 in MRSA biofilm formation and dispersal. Biofilm was quantitated in microplates using a crystal violet binding assay. Extracellular DNA (eDNA) was isolated from colony biofilms and analyzed by agarose gel electrophoresis. In some experiments, broth or agar was supplemented with sub-MIC amoxicillin to induce biofilm formation. Biofilm erosion was quantitated by culturing biofilms on rods, transferring the rods to fresh broth, and enumerating CFUs that detached from the rods. Biofilm sloughing was investigated by culturing biofilms in glass tubes perfused with broth and measuring the sizes of the detached cell aggregates. We found that a *nuc1* mutant strain produced significantly more biofilm and more eDNA than a wild-type strain in both the absence and presence of sub-MIC amoxicillin. *nuc1* mutant biofilms grown on rods detached significantly less than wild-type biofilms. Detachment was restored by exogenous DNase or a wild-type *nuc1* gene on a plasmid. In the sloughing assay, *nuc1* mutant biofilms released cell aggregates that were significantly larger than those released by wild-type biofilms. Our results suggest that Nuc1 modulates biofilm formation, biofilm detachment, and the sizes of detached cell aggregates. These processes may play a role in the spread and subsequent survival of MRSA biofilms during biofilm-related infections.

## INTRODUCTION

*Staphylococcus aureus* is a major public health burden in both community and hospital settings (Percival *et al*., 2015). In hospitals, *S. aureus* is a major cause of infections associated with implanted medical devices because of the ability of most *S. aureus* strains to form biofilms on biomaterial surfaces (Schilchera & Horswill, 2020). *S. aureus* biofilms on implanted devices can act as a nidus of infection when cells detach from the biofilm and seed other parts of the body such as the blood, lung, or bladder (Archer *et al*., 2011). In addition, nearly half of all *S. aureus* hospital isolates are methicillin-resistant *S. aureus* (MRSA), a mul-drug-resistant strain that further complicates treatment of these common nosocomial infections (Diekema *et al*., 2019).

*S. aureus* secretes numerous virulence factors into its surrounding environment that contribute to host colonization and virulence. These include extracellular enzymes, pore-forming toxins and superantigens (Busche *et al*., 2018). Among these is micrococcal nuclease (Nuc1), also known as thermonuclease, a secreted, Ca+2-dependent, thermostable phosphodiesterase that hydrolyzes both RNA and DNA (Olson *et al*., 2013). Nuc1 is produced by all strains of *S. aureus. nuc1* mutant strains exhibit decreased survival in mouse models of peritonitis (Olson *et al*., 2013) and intranasal infection (Berends *et al*., 2010). Nuc1 may contribute to virulence in vivo by facilitating escape from neutrophil extracellular traps (Berends *et al*., 2010; Bhattacharya *et al*., 2020; Thammavongsa *et al*., 2013). Nuc1, along with other exoenzymes such as proteases, lipases, and hyaluronidases, may also contribute to tissue invasion or nutrient acquisition in vivo (Tam & Torres, 2019). All strains of *S. aureus* also produce a second extracellular nuclease termed Nuc2 (Tang *et al*., 2008; Kiedrowski *et al*., 2014). Like Nuc1, the homologous Nuc2 enzyme is Ca+2-dependent, thermostable, and able to hydrolyze both DNA and RNA (Tang *et al*., 2008; Kiedrowski *et al*., 2014). Nuc2 is expressed in vivo, and purified Nuc2 protein inhibits *S. aureus* biofilm formation in microtiter plate wells and detaches preformed biofilms (Kiedrowski *et al*., 2014), suggesting that Nuc2 may also play a role in biofilm formation and dispersal. However, the *nuc1* and *nuc2* genes are located in different regions of the *S. aureus* chromosome, and the Nuc2 protein remains bound to the cell surface (Tang *et al*., 2008; Kiedrowski *et al*., 2014).

Extracellular DNA (eDNA) is the most common component of biofilms produced by most MRSA strains (Sugimoto *et al*., 2018). eDNA consists of genomic DNA released by lysis of a subpopulation of cells within the biofilm (Archer *et al*., 2011; Mann *et al*., 2009). eDNA may help stabilize biofilm matrix by binding to secreted eDNA-binding proteins and membrane-attached lipoproteins (Kavanaugh *et al*., 2019) or to extracellular poly-N-acetylglucosamine polysaccharide (Mlynek *et al*., 2020). Previous studies showed that biofilms produced by *nuc1* mutant strains contain more eDNA than wild-type biofilms (Kiedrowski *et al*., 2011), and that biofilm formation is enhanced in *nuc1* mutant strains (Mann *et al*., 2009; Kiedrowski *et al*., 2011). In addition, expression of *nuc1* is repressed during biofilm formation (Olson *et al*., 2013; Kiedrowski *et al*., 2011). These findings suggest that exogenous Nuc1 can degrade eDNA, thereby decreasing the adhesive properties of the biofilm matrix.

In the present study we investigated the role of Nuc1 and Nuc2 in biofilm formation using MRSA strain LAC, a USA300 clone. By comparing a wild-type strain to *nuc1*-, *nuc2*-, and *nuc1*-/*nuc2*- mutant strains, we found that Nuc1, but not Nuc2, modulates the amount of eDNA in the biofilm matrix, the adhesiveness of the biofilms, and the amount of biofilm induced by low-dose amoxicillin. We further investigated the role of Nuc1 in two different mechanisms of biofilm dispersal, namely, erosion and sloughing. Biofilm erosion refers to the release of single cells or small clusters of cells from a biofilm, whereas sloughing refers to the detachment of large portions of the biofilm (Kaplan, 2010). Results from assays comparing biofilm dispersal in wild-type and *nuc1* mutant strains suggests that Nuc1 not only mediates detachment of cells from MRSA biofilms, but also modulates the sizes of the detached cell aggregates.

## MATERIALS AND METHODS

### MATERIALS AND METHODS

#### Bacterial strains, media, and growth conditions

The bacterial strains used in this study are listed in Table 1. Bacteria were cultured on Tryptic Soy agar plates or in filter-sterilized Tryp-c Soy broth (BD Diagnostics). All cultures were incubated at 37°C. For plasmid-harboring strains, media were supplemented with 10 μg/ml chloramphenicol. To induce biofilm formation, media were supplemented with sub-MIC concentrations of amoxicillin ranging from 0.002-2 μg/ml. To induce biofilm detachment, broth was supplemented with 10 μg/ml recombinant human DNase I (Genentech), which was previously shown to efficiently detach *S. aureus* biofilms from polystyrene microtiter plate wells (Kaplan *et al*., 2012). Bacterial inocula were prepared in sterile broth from 24-h-old agar colonies and then passed through a 5-μm pore-size syringe filter to enrich for single cells as previously described (Izano *et al*., 2008).

**Table 1.**
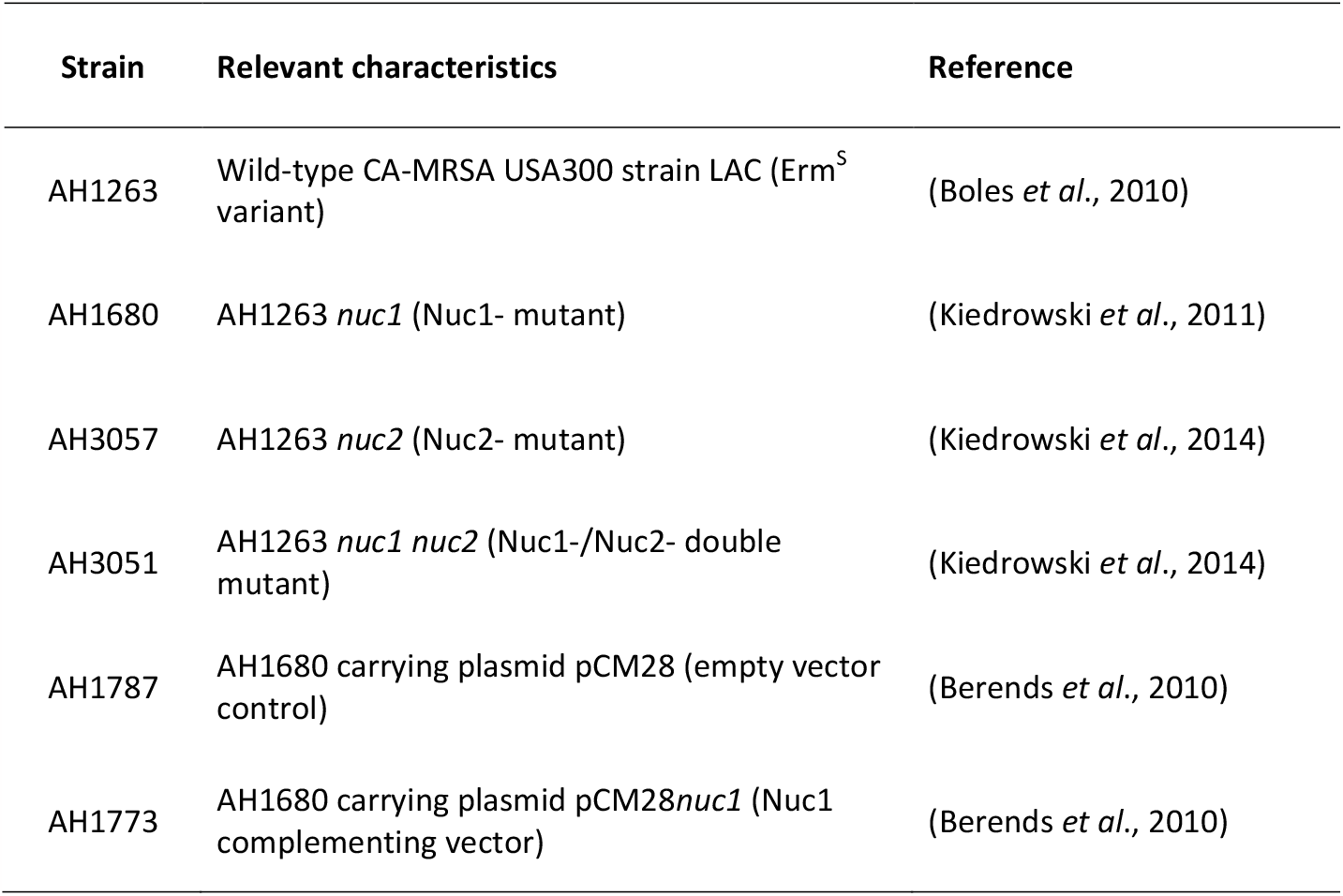
*S. aureus* strains used in this work.

| Strain | Relevant characteristics | Reference |
| --- | --- | --- |
| AH1263 | Wild-type CA-MRSA USA300 strain LAC (Erm<sup>S</sup> variant) | (Boles *et al.*, 2010) |
| AH1680 | AH1263 *nuc1* (Nuc1- mutant) | (Kiedrowski *et al.*, 2011) |
| AH3057 | AH1263 *nuc2* (Nuc2- mutant) | (Kiedrowski *et al.*, 2014) |
| AH3051 | AH1263 *nuc1 nuc2* (Nuc1-/Nuc2- double mutant) | (Kiedrowski *et al.*, 2014) |
| AH1787 | AH1680 carrying plasmid pCM28 (empty vector control) | (Berends *et al.*, 2010) |
| AH1773 | AH1680 carrying plasmid pCM28*nuc1* (Nuc1 complementing vector) | (Berends *et al.*, 2010) |

#### 6-well microtiter plate biofilm assay

Aliquots of inocula (4-ml each; ca. 102 CFU/ml) were transferred to the wells of a 6-well microtiter plate (Falcon #353046), and the plate was incubated statically under conditions of low vibration as previously described (Kaplan & Fine, 2002). After 18 h, a 1-cm2 area in the center of each well was photographed. Wells were then rinsed with water and stained for 1 min with 4 ml of Gram’s crystal violet. Wells were rinsed with water and dried, and the same area of each well was rephotographed.

#### 96-well microtiter plate biofilm assay

Aliquots of inocula (180 μl each; 105-106 CFU/ml) were transferred to the wells of a 96-well microtiter plate (Corning #3599) containing 20 μl of antibiotic dissolved in water at a concentration equal to 10 times the desired final concentration. Control wells were filled with 180 μl of inoculum and 20 μl of water, or 180 of sterile broth and 20 μl of water. The plates were incubated for 18 h. The amount of biofilm biomass in each well was quantitated by rinsing the wells with water and staining for 1 min with 200 μl of Gram’s crystal violet. The wells were then rinsed with water and dried. The bound crystal violet was dissolved in 200 μl of 33% acetic acid and quantitated by measuring the absorbance of the wells at 595 nm.

#### Isola&on and analysis of extracellular DNA

Extra-cellular DNA was isolated from lawns of bacteria cultured on agar, also known as colony biofilms, using the method described by Karwacki *et al*. (2013). Briefly, 100-μl aliquots of inoculum (>108 CFU) were spread onto agar plates supplemented with 0 or 0.2 μg/ml amoxicillin. After incubation for 24 h, the cell paste from each plate was transferred to a separate, pre-weighed 1.5-ml microcentrifuge tube, weighed, and resuspended in TE buffer at a concentration of 1 mg/ml. The tube was mixed by vortex agitation for 10 min, and the cells were pelleted by centrifugation. The supernatant was sterilized by passage through a 0.2-μm-pore-size filter. A 20-μl volume of colony biofilm extract was analyzed by agarose gel electrophoresis, and the DNA was visualized by staining with ethidium bromide.

#### Biofilm erosion assay

An apparatus to measure biofilm erosion under static conditions (Fig. 1A) was constructed and employed as follows. First, a 2-mm diam hole was drilled in the center of a 35-mm diam Petri dish lid (Sarstedt # 83.1800.001). Next, a 100-mm polystyrene or glass rod was placed in the hole and secured in a vertical position with hot-melt adhesive. The polystyrene rods (1.5-mm diam) were purchased from Plastruct (City of Industry CA), and the glass rods were made from a 1.9-mm diam open-one-end capillary tubes (Stuart Glass # EW-03013-64) with the closed end facing down. The rods were positioned so that 5 mm protruded from the top of the lid (Fig. 1A). The apparatus were sterilized with 70% ethanol, air dried, and then placed on top of 50-ml conical centrifuge tubes (Corning # 430290) containing 20 ml of bacterial inoculum at 105-106 CFU/ml. Approximately 30 mm of the rod or capillary tube was suspended in the broth. After incubation for 16 h, the apparatus were carefully transferred to fresh tubes containing 25 ml of pre-warmed sterile broth and incubated for 1 min to remove loosely adherent cells. The apparatus were then transferred to fresh tubes containing 25 ml of pre-warmed sterile broth. After 2 h, the apparatus were removed from the centrifuge tubes. To enumerate the CFUs in the broth, the broth was mixed by vortex agitation for 30 s, serially diluted in phosphate buffered saline (PBS), and then plated on agar for CFU enumeration. To enumerate the CFUs on the rods, the rods were detached from the Petri dish lids, placed in 15-ml conical centrifuge tubes containing 10 ml of PBS, and sonicated for 30 s at 40% duty cycle in an ultrasonic homogenizer equipped with a 1/8 in. probe (Fisher Scientific model 505). Sonicates were serially diluted in PBS and plated on agar for CFU enumeration. The fraction of detached cells (percent CFUs in broth) was calculated by dividing the total number of CFUs in the broth by the total number of CFUs in the broth plus the total number of CFUs on the rod. Assays were performed in duplicate to quadruplicate tubes for each experimental condition.

**Fig. 1.**
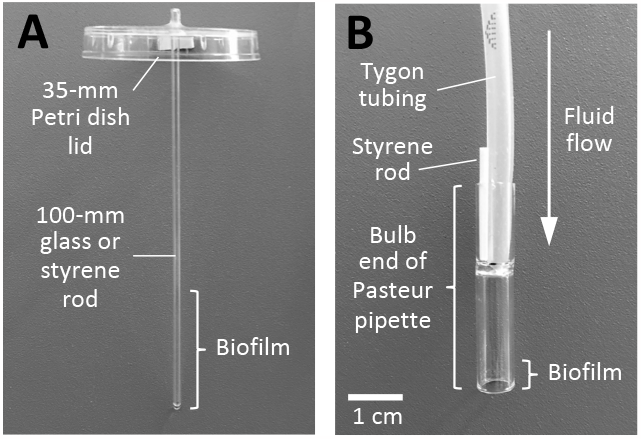
Apparatus used to measure biofilm erosion (A) and biofilm sloughing (B). The construction and use of these apparatus are presented in the Methods section.

#### Biofilm sloughing assay

An apparatus to measure biofilm sloughing under flow conditions (Fig. 1B) was constructed and employed as follows. First, biofilm reactors were constructed by cutting 38-mm-long sections from the bulb ends of standard 9-in. flint glass Pasteur pipettes (VWR # 14672-380) using a glass cutter. The glass reactors were rinsed in 1% HCl for 2 h, rinsed with 70% ethanol and ultrapure water, and then air dried. Glass reactors were inoculated by standing them (bulb-side-down) in a 6-well microtiter plate well containing 2 ml of bacterial inoculum at 105-106 CFU/ml, which brought the bottom 5-mm of each reactor in contact with the inoculum. After 10 min, the reactors were connected to 1/8 × 3/16 in. silicon tubing (Tygon 3350 #ABW00006) and perfused with fresh broth for 48 h at a rate of 5-7 ml/h. A short 1.5-mm-diam polystyrene rod (Fig. 1B) functioned to secure the tubing to the glass reactor and to form a vent which allowed the reactor to act as a continuous drip chamber. A total of four reactors were perfused in each experiment. To enumerate CFUs in the eluate, drops were collected directly into 1.5-ml microcentrifuge tubes, mixed by vortex agitation for 30 s, serially diluted in PBS, and plated on agar for CFU enumeration. To measure the sizes of the detached cell aggregates, drops were collected directly onto glass microscope slides and then fixed, stained with Gram’s crystal violet, and photographed at 400× under an inverted microscope. For each strain, the distribution of particle sizes in five random microscopic fields was analyzed using ImageJ software. To visualize detached cell aggregates directly, drops of eluate were collected into the wells of a 6-well microtiter plate and then photographed on a dark background.

#### Statistics and reproducibility of results

Biofilm erosion assays were performed in triplicate tubes and were repeated 2-6 times. The significance of differences between percent CFUs in broth values was calculated using a 2-tailed Student’s t-test for pairwise comparisons, and a one-way ANOVA with Tukey’s post hoc analysis for comparison of more than two groups. A P value of <0.05 was considered significant. Biofilm erosion assays were performed in duplicate reactors and were repeated twice. The significance of differences between particle size distributions in the sloughing assay was calculated using a two-way Chi squared test.

## RESULTS

### Biofilm formation by wild-type and nuclease mutant strains in 6-well microtiter plate wells

Wild-type, *nuc1*-, *nuc2*- and *nuc1*-/*nuc2*- strains were cultured in broth in 6-well microtiter plates at low densities and under conditions of low vibration. After 18 h, the wells were photographed (Fig. 2, top panels). All four strains produced surface attached biofilms under these conditions. *nuc1*- and *nuc1*-/*nuc2*- biofilms appeared denser and more compact than those of the wild-type and *nuc2*- strains. After rinsing the wells with water and staining with crystal violet, biofilms of the *nuc1*- and *nuc1*-/*nuc2*- strains remained firmly attached to the surface, whereas biofilms of the wild-type and *nuc2*- strains were readily detached (Fig. 2, bottom panels). These results suggest that Nuc1, but not Nuc2, contributes to USA300 biofilm formation under these conditions.

**Fig. 2.**
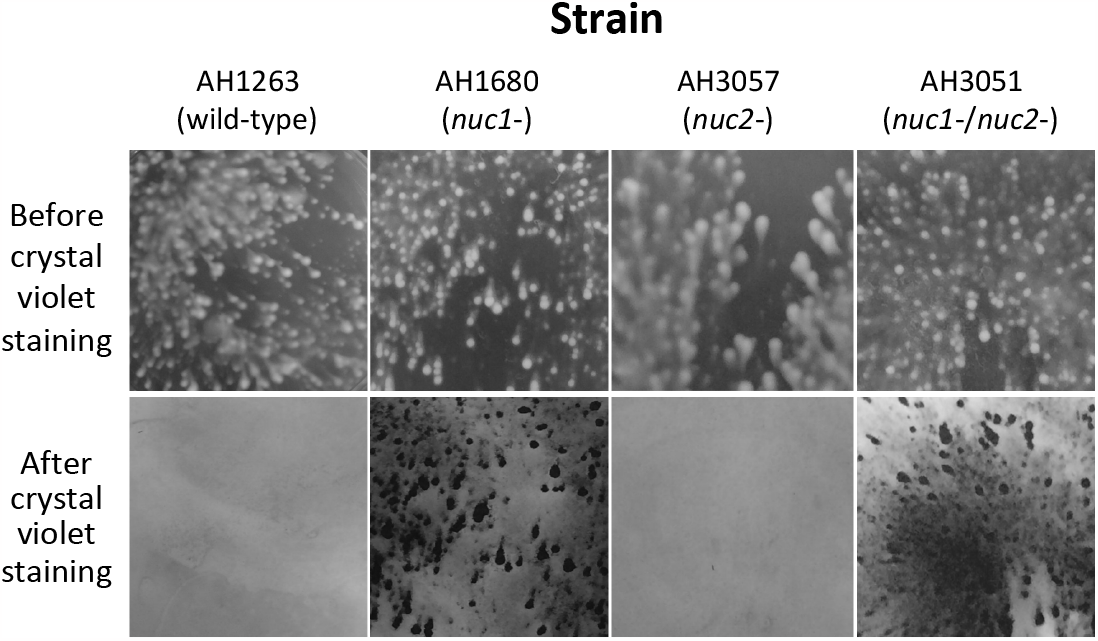
Biofilm formation by *S. aureus* wild-type and nuclease mutant strains in 6-well tissue culture-treated microplates under conditions of low vibration. Top panels show surface-attached growth after incubation for 16 h. Bottom panels show the identical areas of the plate after they were rinsed with water and stained with crystal violet. The size of each imaged area is 1 cm^2^.

### Induction of biofilm formation by sub-MIC amoxicillin in wild-type and nuclease mutant strains

Previous studies showed that low doses of amoxicillin induce biofilm in *S. aureus* USA300 by a mechanism that involved autolysis and eDNA release (Kaplan *et al*., 2012; Mlynek *et al*., 2016). To determine whether Nuc1 or Nuc2 play a role in this process, we used a 96-well crystal violet binding assay to quantitate biofilm formation by wild-type, *nuc1*-, *nuc2*- and *nuc1*-/*nuc2*- strains cultured in sub-MIC amoxicillin concentrations ranging from 0.002-2 μg/ml (Fig. 3). The MIC of amoxicillin against strain USA300 is >8 μg/ml (Kaplan *et al*., 2012). Low dose amoxicillin induced biofilm formation in all four strains, with maximum induction occurring at a concentration of 0.2 μg/ml. However, biofilm induction occurred over a greater range of amoxicillin concentrations and resulted in a significantly greater amount of biofilm biomass in the *nuc1*- and *nuc1*-/*nuc2*- strains compared to the wild-type and *nuc2*- strains (Fig. 3). These results suggest that Nuc1, but not Nuc2, can modulate the structure of MRSA biofilms under both antibiotic-induced and uninduced conditions. These results are consistent with those of previous studies showing that nuclease activity is not required for ⍰-lactam-induced MRSA biofilm formation (Kaplan *et al*., 2012).

**Fig. 3.**
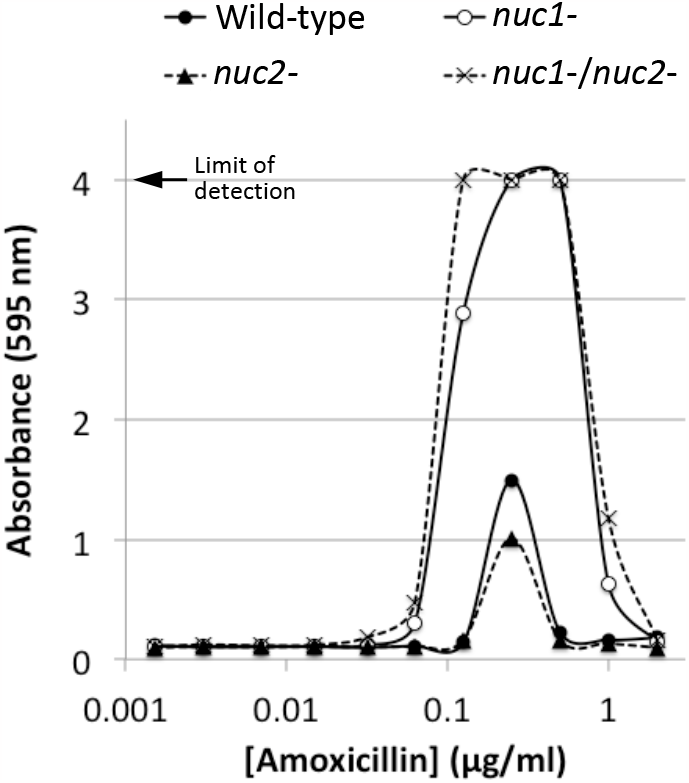
Biofilm formation by *S. aureus* wild-type and nuclease mutant strains in 96-well microtiter plates in the presence of increasing concentrations of sub-MIC amoxicillin. Absorbance at 595 nm is proportional to biofilm biomass.

### eDNA production in wild-type and nuclease mutant strains

To investigate the role of Nuc1 and Nuc2 in USA300 eDNA production, we isolated eDNA from colony biofilms of wild-type, *nuc1*-, *nuc2*-, and *nuc1*-/*nuc2*- strains and analyzed the DNA by agarose gel electrophoresis (Fig. 4A). The *nuc1*- and *nuc1*-/*nuc2*- strains produced significantly more high molecular weight eDNA (> 25 kb) than the wild-type and *nuc2*- strains. All four strains produced high molecular weight DNA in the presence of low-dose amoxicillin (Fig. 4B), which suggests that Nuc2 may regulate eDNA production under antibiotic-induced conditions. Taken together, our results demonstrate that high molecular weight eDNA production inversely correlates with Nuc1 production (Fig. 4), but positively correlates with biofilm tenacity (Fig. 2) and antibiotic-induced biofilm (Fig. 3).

**Fig. 4.**
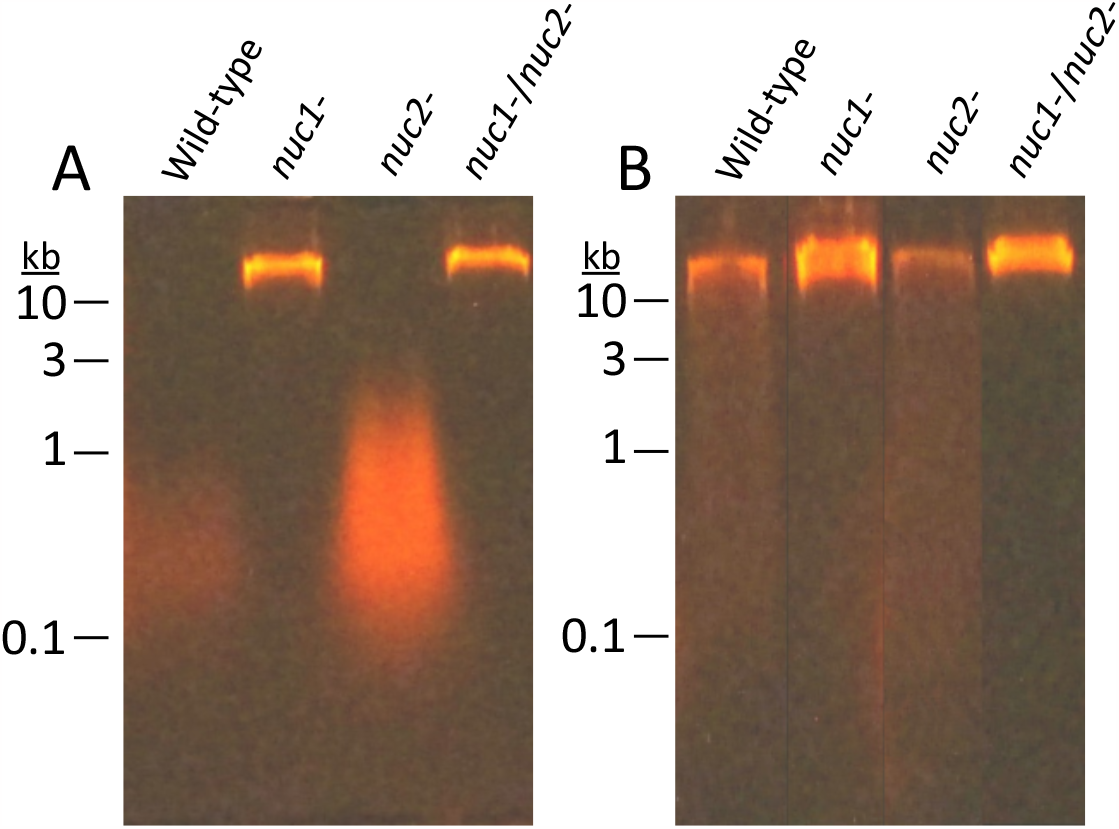
Agarose gel electrophoretic analysis of extracellular DNA (eDNA) produced by *S. aureus* wild-type and nuclease mutant strains. eDNA was harvested from colony biofilms cultured on TSA (A) or TSA supplemented with 0.2 μg/ml amoxicillin (B). Numbers at the left indicate the sizes of molecular weight markers electrophoresed in an adjacent lane.

### Role of Nuc1 in biofilm erosion

We next investigated the role of Nuc1 in MRSA biofilm erosion. We focused on Nuc1 because Nuc1 appears to be the primary modulator of USA300 eDNA production and biofilm formation under uninduced conditions. Biofilm erosion was quantitated by culturing wild-type and *nuc1*- biofilms on polystyrene or glass rods, transferring the rods to fresh broth, and enumerating the CFUs that detached from the rods after 2 h (Fig. 1A). We calculated the percentage of CFUs in the broth by dividing the number of CFUs in the broth by the number of CFUs in the broth plus the number of CFUs that remained on the rod. The total CFUs ranged from 108-109 for both polystyrene and glass rods. Fig. 5A shows the results from multiple biofilm erosion assays performed on polystyrene rods. On average, significantly more CFUs detached from wild-type biofilms compared to the number that detached from *nuc1*- mutant biofilms (68% of CFUs in broth for wild-type versus 38% for *nuc1*- mutant). The same erosion phenotype was observed when biofilms were cultured on glass rods (76% CFUs in broth for wild-type versus 38% for *nuc1*-) (Fig. 5B). Greater than 90% of the CFUs were present in the broth when *nuc1*- mutant biofilms were allowed to erode in broth supplemented with DNase (Fig. 5B), or when the *nuc1*- strain carried a wild-type *nuc1* gene on plasmid. (Fig. 5C). These findings confirm that Nuc1 mediates MRSA biofilm erosion.

**Fig. 5.**
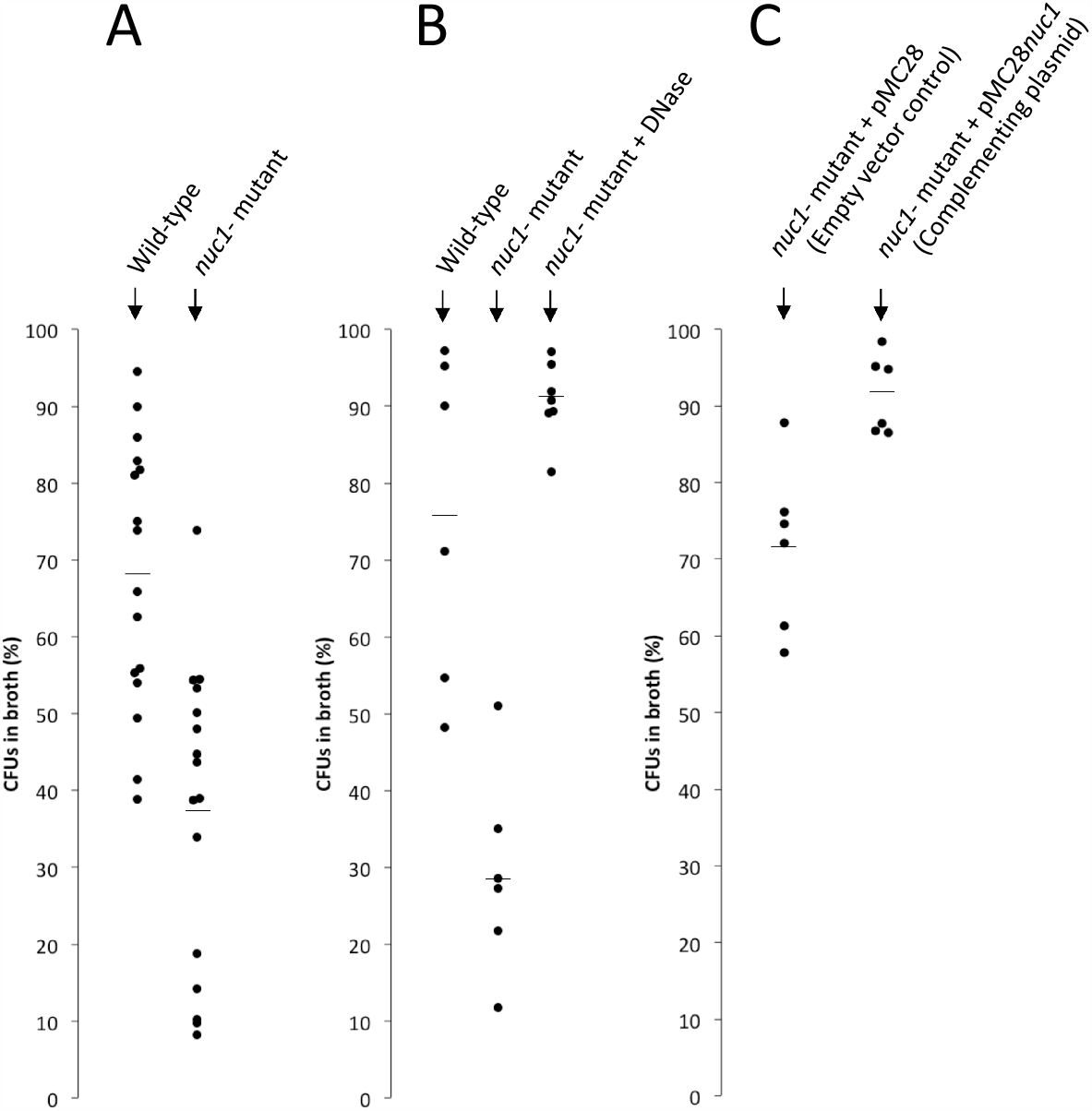
Erosion of *S. aureus* wild-type and *nuc1-* mutant biofilms from polystyrene rods (A) and glass capillary tubes (B and C). Values show the percent of total CFUs in the broth after 2 h. Each dot represents one rod. (A) Wild-type and *nuc1*- mutant. (B) DNase I causes erosion in the *nuc1*- mutant. (C) A *nuc1* complementing plasmid (pCM28*nuc1*) induces erosion in the *nuc1*- mutant.

### Role of Nuc1 in biofilm sloughing

Biofilm sloughing is the detachment of large portions of a biofilm, which usually occurs in the later stages of biofilm formation (Kaplan, 2010). To quantitate biofilm sloughing, we grew wild-type and *nuc1*- mutant biofilms in glass tubes perfused with fresh broth and measured the number of CFUs and sizes of detached cell aggregates in the eluate (Fig. 1B). Both strains formed biofilms in the glass tube reactors and dispersed. Over the course of the 48-h assay, the *nuc1*- mutant biofilm grew inside the silicone tubing that supplied fresh broth to the reactor as evidenced by the presence of crystal violet staining material inside the tube (Fig. 6A). Biofilms of both strains dispersed at high levels from 18-52 h (Fig. 6B), but *nuc1*- mutant biofilms released at least ten times more CFUs than wild-type biofilms at all ‘me points tested, probably because of a larger amount of biofilm biomass. We also measured the sizes of particles that detached from wild-type and *nuc1*- mutant biofilms using ImageJ software. Graphs showing the distribution of particle sizes from two independent sloughing experiments are shown in Figs. 6C-D. Characteris&cs of the detached par&cles from both experiments are summarized in Table 2. In both experiments, the distributions of particle sizes were significantly different (P < 0.0001; two-way Chi squared test) (Figs. 6C-D). In addition, the average number of particles in the *nuc1*- eluate was significantly less than the number in the wild-type eluate, and the average particle size in the *nuc1*- eluate was significantly greater than that in the wild-type eluate in both experiments (Table 2). After 48-52 h of perfusion, macroscopic clusters of cells were present in the *nuc1*- eluate, but not in the wild-type eluate (Fig. 6E).

**Table 2.** Characteristics of cell aggregates in the eluate of wild-type and *nuc1*- biofilms perfused with broth for 20-22 h. Results from two independent experiments are shown.

|  | EXPT. 1 (20 h) |  | EXPT. 2 (22 h) |  |
| --- | --- | --- | --- | --- |
|  | AH1263<br>(Wild-type) | AH1680<br>(*nuc1*-) | AH1263<br>(Wild-type) | AH1680<br>(*nuc1*-) |
| Average number of particles (± sd) per microscopic field | 1,436<br>± 438 | 135<br>± 33* | 1,072<br>± 287 | 492<br>± 108* |
| Average particle size ($\mu\text{m}^2$) | 15 | 144* | 317 | 723* |

\* Significantly different from strain AH1263 ( $P < 0.01$ , 2-tailed Student's *t*-test).

**Fig. 6.**
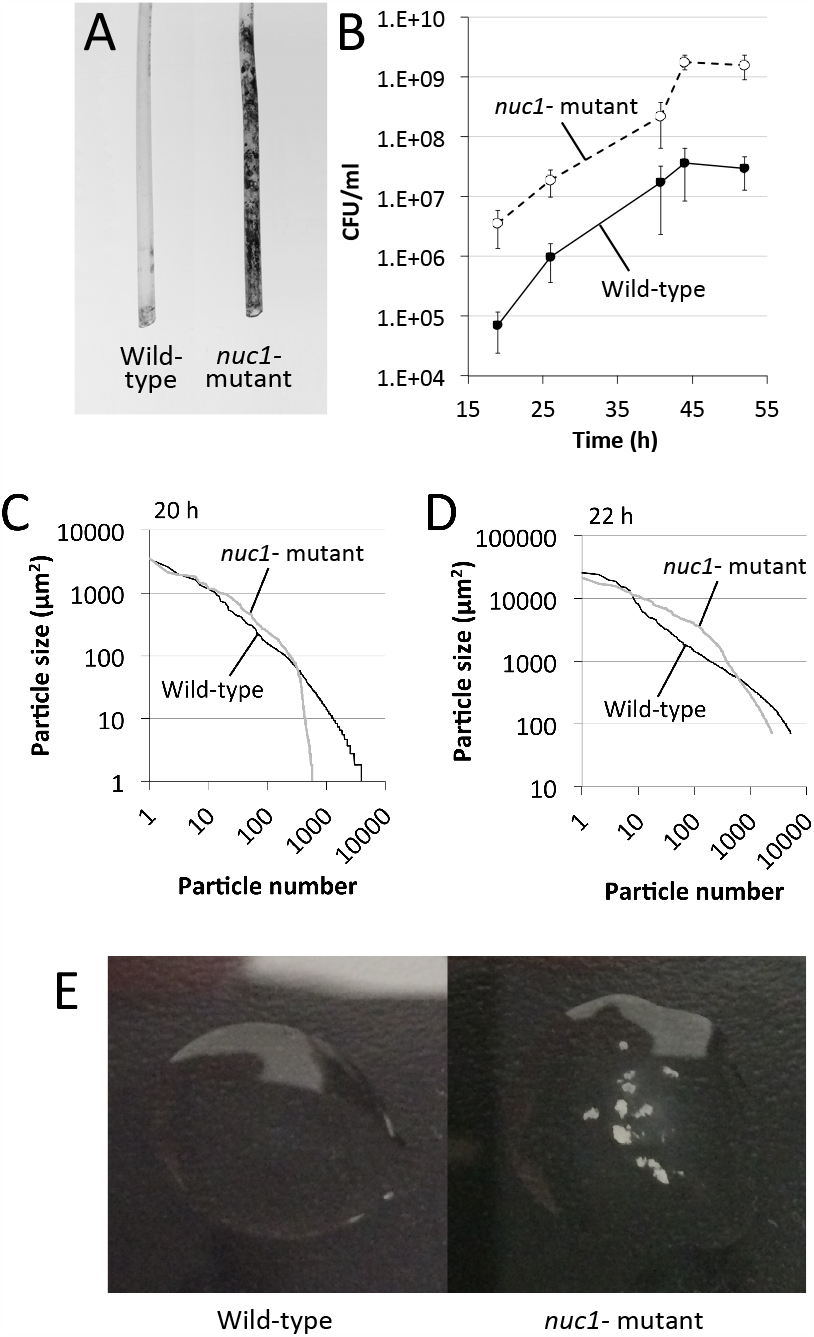
Biofilm sloughing assay. (A) Photographs of the sections of silicone tubing located directly above the glass biofilm reactors. The tubing sections were stained with crystal violet after 48 h of perfusion. (B) CFU/ml in the flowthrough from 17-52 h. (C and D) Distribution of particle sizes in the eluates from two independent sloughing experiments. Eluates were analyzed after 20 h (panel C) or 22 h (panel D) of perfusion. (E) Photographs of drops of eluate after 48 h of perfusion.

## DISCUSSION

Previous studies showed that biofilms produced by *S. aureus nuc1* mutant strains are thicker and contain more eDNA than wild-type biofilms (Mann *et al*., 2009; Kiedrowski *et al*., 2011; Forson 2022), and that *nuc1* expression is repressed during biofilm formation (Olson *et al*., 2013; Kiedrowski *et al*., 2011). These findings are consistent with the hypothesis that Nuc1 degrades eDNA and decreases the adhesive properties of the biofilm matrix. A corollary hypothesis is that Nuc1 may function as an endogenous mediator of biofilm dispersal in this species. Dispersal of biofilms from indwelling medical devices is a clinically relevant process that may lead to invasive infections such as ventilator-associated pneumonia and endocarditis (Adair *et al*., 1999). Although Moormeier *et al*. (2014) showed that Nuc1 mediates the detachment and dispersal of cells prior to the development of characteristic tower-like structures in the early stages of *S. aureus* biofilm formation, no studies have directly tested the hypothesis that Nuc1 mediates detachment and dispersal of cells from mature *S. aureus* biofilm.

In the present study we constructed apparatus to measure *S. aureus* biofilm erosion and biofilm sloughing. Erosion refers to the continuous release of single cells or small clusters of cells from a biofilm at low levels over the course of biofilm formation, whereas sloughing refers to the sudden detachment of large portions of the biofilm, usually during the later stages of biofilm formation (Kaplan, 2010). We measured biofilm erosion and sloughing in a wild-type MRSA strain and an isogenic *nuc1* mutant strain. We focused on Nuc1 because Nuc2 appears to play a minor role in *S. aureus* biofilm formation (Figs. 3-4) and eDNA release (Fig. 5) under the conditions tested.

To measure biofilm erosion, we cultured biofilms overnight on glass or polystyrene rods, rinsed the rods to remove loosely adherent cells, and then transferred the rods to tubes containing fresh broth. After 2 h, the rods were removed from the tubes and the number of CFUs on the rod and in the broth were measured. Percent biofilm dispersal was calculated as (broth CFUs)/(broth CFUs + rod CFUs) × 100. *nuc1* mutant biofilms detached significantly less than wild-type biofilms (38% versus 68% from polystyrene rods and 28% versus 76% from glass rods; Fig. 6A-B). Detachment by the *nuc1* mutant was restored to wild-type levels by exogenous DNase (Fig. 6B) or a wild-type *nuc1* gene on a plasmid (Fig. 6C), confirming that Nuc1 mediates biofilm erosion under the conditions tested.

To study biofilm sloughing, biofilms were cultured on the bulb ends of glass Pasteur pipettes perfused with fresh broth, and the quantity and sizes of detached cell aggregates in the flow-through were measured over 48 h. Under these conditions, the *nuc1* mutant strain produced significantly more biofilm than the wild-type strain as evidenced by a greater amount of crystal violet-stained material in the silicone tubes leading into the reactors (Fig. 7A), and a significantly greater number of CFUs in the eluate over 5me (Fig. 7B). *nuc1* mutant biofilms released cell aggregates that were significantly larger than those released by wild-type biofilms (Fig. 7C-D and Table 2) as well as visible cell aggregates that were not present in the eluate of wild-type biofilms (Fig. 7E). The sizes of the detached cell aggregates in our study were similar to the sizes of aggregates that detached from *S. aureus* biofilms cultured in glass tubes perfused with brain heart infusion medium (Fux *et al*., 2004) and in silicone tubing perfused with human plasma (Grønnemose *et al*., 2017). Large, detached aggregates of *S. aureus* cells were previously shown to be highly resistant to antibiotics (Fux *et al*., 2004, Haaber *et al*., 2012) and phagocytosis (Alhede *et al*., 2020, Pettygrove *et al*., 2021), and capable of initiating colonization of endothelial cell layers under flow (Grønnemose *et al*., 2017), which suggests that biofilm sloughing may contribute to metastatic infections associated with *S. aureus* (Fux *et al*., 2004).

Our results demonstrate that Nuc1 modulates biofilm erosion and sloughing in *S. aureus* USA300. A homologous nuclease was shown to modulate eDNA production, biofilm dispersal, and biofilm aggregate size in nontypeable Haemophilus influenzae (Cho *et al*., 2015). Since eDNA is a structural component of Pseudomonas aeruginosa biofilms (Allesen-Holm *et al*., 2006), it is possible that Nuc1 may passively contribute to P. aeruginosa biofilm formation and dispersal during *S. aureus*/P. aeruginosa coinfection in the airways of cystic fibrosis patients (Wieneke *et al*., 2021).

## ACKNOWLEDGEMENTS

We thank Gabrielle Kyle-Lion, Roxanna Stapleton, and Amelia Crabtree (American University) for technical assistance.

